# scDecouple: Decoupling cellular response from infected proportion bias in scCRISPR-seq

**DOI:** 10.1101/2023.01.31.526445

**Authors:** Qiuchen Meng, Lei Wei, Kun Ma, Ming Shi, Xinyi Lin, Joshua W. K. Ho, Yinqing Li, Xuegong Zhang

**Affiliations:** MOE Key Lab of Bioinformatics & Bioinformatics Division BRNIST, Department of Automation, Tsinghua University, Beijing 100084, China; MOE Key Laboratory of Bioinformatics, Tsinghua University, Beijing, 100084, China; IDG-McGovern Institute for Brain Research, Center for Synthetic and Systems Biology, School of Pharmaceutical Sciences, Tsinghua University, Beijing, 100084, China; School of Biomedical Sciences, Li Ka Shing Faculty of Medicine, The University of Hong Kong, Pokfulam, Hong Kong SAR, China; Laboratory of Data Discovery for Health Limited (D24H), Hong Kong Science Park, Hong Kong SAR, China; Center for Synthetic and Systems Biology, School of Life Sciences and School of Medicine, Tsinghua University, Beijing 100084, China

## Abstract

**Motivation:** scCRISPR-seq is an emerging high-throughput CRISPR screening technology that combines CRISPR screening with single-cell sequencing technologies. It provides rich information on the mechanism of gene regulation. However, when scCRISPR-seq is applied in a population of heterogeneous cells, the true cellular response to perturbation is coupled with infected proportion bias of guide RNAs (gRNAs) across different cell clusters. The mixing of these effects introduces noise into scCRISPR-seq data analysis and thus obstacles to relevant studies.

**Results:** We developed scDecouple to decouple true cellular response of perturbation from the influence of infected proportion bias. scDecouple first models the distribution of gene expression profiles in perturbed cells and then iteratively finds the maximum likelihood of cell cluster proportions as well as the cellular response for each gRNA. We demonstrated its performance in a series of simulation experiments. By applying scDecouple to real scCRISPR-seq data, we found that scDecouple enhances the identification of biologically perturbation-related genes. scDecouple can benefit scCRISPR-seq data analysis, especially in the case of heterogeneous samples or complex gRNA libraries.

**Availability:** scDecouple is freely available at https://github.com/MengQiuchen/scDecouple.

## 1 Introduction

With the advancement in single-cell sequencing and CRISPR (clustered regularly interspaced short palindromic repeats) technologies, scCRISPR-seq has emerged as a novel high-throughput gene function profiling method^1^. scCRISPR-seq first leverages CRISPR to perturb a set of genes and then assesses the resulting profiles of each perturbation by single-cell sequencing. There exist multiple scCRISPR-seq protocols, including Perturb-seq^2–4^, CROP-seq^5^, CRISP-seq^6^, Mosaic-seq^7^, SpearATAC^8^, and CRISPR-sciATAC^9^. In a typical scCRISPR-seq protocol, a pool of guide RNAs (gRNAs) targeting different genes are usually packed into lentivirus and then introduced into cells. These gRNAs each introduce perturbation to a subgroup of cells. Then, single-cell sequencing^10-14^ is used to capture one or more types of profiles for each cell, such as single-cell RNA sequencing (scRNA-seq)^10,11^, single-cell ATAC sequencing (scATAC-seq)^12,13^, and Cellular Indexing of Transcriptomes and Epitopes by Sequencing (CITE-seq)^14^. These scCRISPR-seq methods enable high-throughput perturbations as well as data-rich read-outs for each perturbation, providing informative data for gene regulation study^2,15–17^, disease target identification^4^, and drug development^18^. Obtaining the true effect of each perturbation is a prerequisite for scCRISPR-seq analysis. A primary challenge lies in the inability to directly measure the original expression profile of perturbed cells. In typical experimental settings, a control group, comprising either unperturbed cells or those infected by non-targeting (NT) gRNAs, is introduced to approximate the profiles of cells before perturbations^2,15,19^. For every gene, the cellular response to a specific perturbation is quantified by computing the fold change (FC) in its average expression level between cells subjected to the perturbation (termed as the perturbation group) and the control group. This approach is not accurate enough, especially in experiments with samples comprising multiple cell clusters or extensive gRNA libraries. There exist significant differences in the proportions of cell clusters between the perturbation group and the control group^2,4^. This difference might be due to the inconsistent infection efficiencies of different gRNAs, inherent differences in growth rates among different cell clusters, or the sampling bias in sequencing. This effect causes the average gene expression level of the control group different from the true original expression profile of the perturbation group. We refer to the distortion introduced by this as “infected proportion bias”. With the development of scCRISPR-seq technologies, the increasing heterogeneity of samples^4^ and the expanding complexity of gRNA libraries^15^ intensify the importance of addressing these challenges.

Here, we developed scDecouple as a solution to decouple cellular responses from infected proportion bias by employing maximum likelihood estimation. scDecouple leverages a Gaussian mixture model to depict the distribution of gene expression profiles in the principal component (PC) space. Through the expectation-maximization (EM) algorithm, we iteratively approximate the genuine cluster proportion of infected cells along with their cellular responses. We evaluated the performance of scDecouple on both synthetic and real-world datasets. The generation of synthetic data considered various parameter settings, including the distance between clusters, and the infection ratio. scDecouple consistently exhibited good performance across different parameter settings. Application to real-world scCRISPR-seq datasets revealed that scDecouple not only provided more precise cellular responses but also enhanced biologically relevant pathway identification and gene ranking. scDecouple facilitates a deeper comprehension of perturbation consequences and paves the way for advancing intricate scCRISPR-seq protocols.

### 2 Methods

### 2.1 Mathematical description of infected proportion bias

We mathematically described the entire experimental process of scCRISPR-seq. Without loss of generality, we assumed that there are two clusters in the experimental cells before perturbation (Fig. 1A). Their average expression (***μ**_X_*) is approximately equal to the average expression of cells in the control group (***μ**_Z_*). Due to the infected proportion bias, the average expression of cells may differ from ***μ***_*Z*_ even taking no consideration about the expression perturbation caused by gRNAs. We named this average expression as ***μ**_S_* (Fig. 1B). Besides, we assumed that the perturbation brought by a gRNA introduces the same alteration to expression profiles of all cells in the perturbation group in the feature space, leading the average expression of the perturbation group alters from ***μ**_S_* to the observed one μ_Y_ (Fig. 1C). The observed change between the control and perturbation group is ***μ**_Y_* − ***μ***_*Z*_, which contains the bias caused by infected proportion bias (***μ**_S_* − ***μ***_*Z*_) and the true cellular response to perturbation (***μ**_Y_* − ***μ**_S_*).

**Fig. 1.**
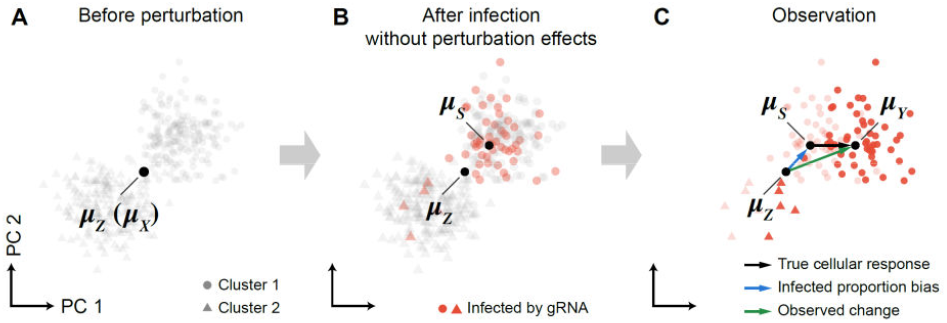
Mathematical description of average expression alteration of the perturbation group. Distributions and average expressions of cells are shown in the feature space. **(A)** Cells before perturbation. **(B)** Cells after infection without the consideration of perturbation effects. **(C)** Cells observed by high-throughput sequencing. Cells in different clusters are represented by different shapes. Cells infected by gRNAs are colored as red.

### 2.2 General workflow of scDecouple

scDecouple comprises four distinct steps: data preprocessing, principal component (PC) selection, decoupling, and downstream analysis (Fig. 2).

**Fig. 2.**
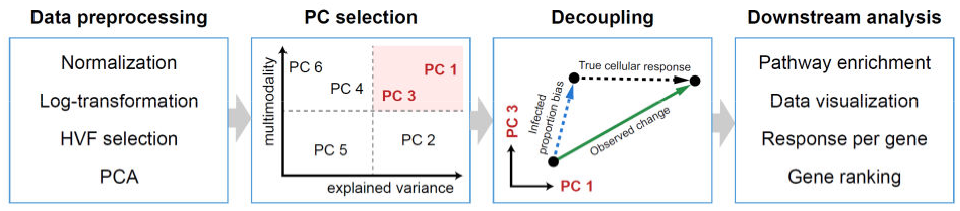
General workflow of scDecouple.

During the data preprocessing phase, the gene expression of each cell is first normalized based on its total expression to reduce the technical variance caused by sequencing depths. The expression then undergoes a log transformation to reduce the effects of extremely high-expressed outliers. After, highly variable features (HVFs) of cells in the control group are selected, and the cell-HVF matrix is transformed into the PC space.

The second step is PC selection. Cells effectively impacted by a gRNA is limited in scCRISPR-seq, making it hard to estimate a mass of parameters. Thus, we designed a PC selection approach to reduce the number of parameters to be estimated. Our analytical derivations revealed that influence of infected proportion bias is most pronounced in PCs with both high variance and multimodality. scDecouple employs this finding as the selection criteria to select PCs bearing infected proportion bias for further analysis. Details of the step are shown in Section 2.2.1.

After, the decoupling procedure is executed on the selected PCs to disentangle cellular response from infected proportion bias. We projected cells in the perturbation group into the PC space defined by HVFs in the control group. We then constructed two Gaussian mixture models (GMMs) for the control and perturbation groups, respectively. Both the models share the same number of cell clusters, but the parameters within these clusters are different. The EM algorithm is used to iteratively estimate the unknown parameters. With this approach, the genuine cluster proportion of infected cells along with their cellular responses are approximated. Details of the step are shown in Section 2.2.2 and 2.2.3.

Finally, we integrated several common downstream analyses based on the decoupled results, including pathway enrichment and data visualization. Besides, the gene expression profile can be regenerated from the PC space, and a ranked list of perturbation-related genes is provided to help explore the cellular response of any gene to a perturbation.

### 2.2.1 PC selection

Despite the large number of sequenced cells, the number of cells effectively impacted by a gRNA is usually small in scCRISPR-seq due to high complexity of gRNA pools and limited gRNA efficiency. For example, the median of cells captured by a single gRNA is 121 in a K562 scCRISPR-seq experiment^15^ and 83 in a Jurkat cell experiment^5^. Reducing the number of parameters to be estimated is crucial to amplify estimation accuracy. To address this, we classified PCs into two groups according to their susceptibility to infected proportion bias. We assumed that the observed gene expression changes can reflect true cellular response in PCs with low influences of infected proportion bias.

We first derived the relationship between the observed change, cellular response and infected proportion bias on each PC. We denoted the expression of cell *i* in the control group on this PC as *z*_*i*_ and the expression of cell j in the perturbation group on this PC as *y*_*j*_. The observed change (OC) is the difference between two groups:

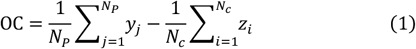

where *N*_*p*_ and *N*_*C*_ represent the number of cells in the perturbation and control group, respectively. We assumed that cells in the control group have *K* clusters, and the average expression profile on this PC and proportion of cluster *k* is *μ*_*k*_ and *δ*_*k*_, respectively. After perturbation, we assumed that the cluster number maintains as K but the proportion of cluster k in the perturbation group changes to λ_k_. There are

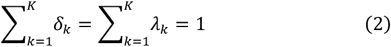

Besides, we assumed that all perturbed cells share a similar response *β*. Thus,

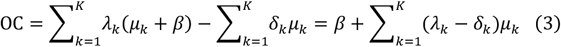

In this equation, the first term represents the cellular response, and second term represents the infected proportion bias.

The infected proportion bias approaches zero under two specific conditions. One condition is that the data on this PC follow a unimodal distribution (*K* = 1, and thus *λ*_l_ = *δ*_l_ = 1). Another condition is that the data variance explained by this PC is low, and thus

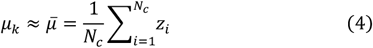

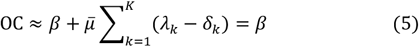

Therefore, in cases where PCs exhibit a unimodal distribution or low explained variance, the observed change is equal or approximately equal to real cellular response. In such scenarios, there is little space for improvement through decoupling. Decoupling is primarily required for PCs that exhibit both multimodality and high explained variance.

The explained variance of a PC is calculated during PC analysis (PCA). The multimodality score of a PC is measured by the Hartigan’s dip test^20^ which quantifies the maximum difference between the empirical distribution and the best-fitting unimodal distribution. PCs that are both multimodal and with high explained variance are selected for decoupling. For any remaining PC, OC calculated by Eq. 1 is directly regarded as the cellular response.

### 2.2.2 Decoupling with GMMs

We used GMMs to describe the gene expression of cells in the control or perturbation group on all selected PCs. The density function of the control group is

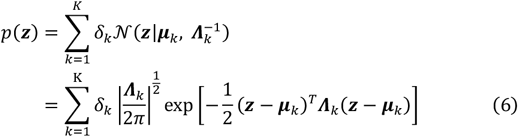

Here, ***μ***_k_ and ***Λ***_k_ represent the expectation (mean) and precision (inverse of variance) of cell cluster k in the control group on all selected PCs, respectively.

The density function of the perturbation group is

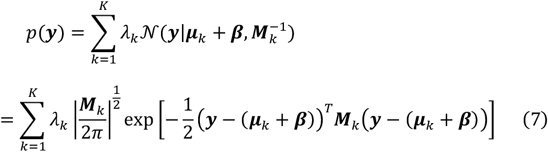

Here, ***M***_k_ represents the and precision of cell cluster *k* in the perturbation group, and ***β*** represents the cellular response on all selected PCs.

### 2.2.3 Inferring cellular response with the EM algorithm

The likelihood function for all observed data is

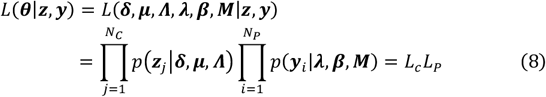

The first component represents the likelihood of the control group (*L*_*c*_), and the second represents the likelihood of the perturbation group (*L*_*p*_).

The EM algorithm is first employed to maximize *L*_*c*_ and estimate parameters ***δ***,***μ, Λ*:**

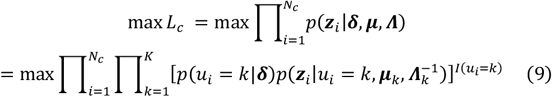

The EM algorithm is then used to maximize *L*_*p*_:

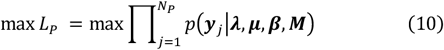

As **μ** has already been estimated in the previous step, the iterative EM algorithm is used to estimate ***λ***,***β***,***M*** .

For the expectation step (E-step):

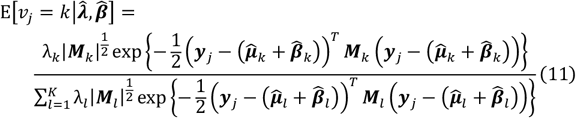

For the maximum step (M-step):

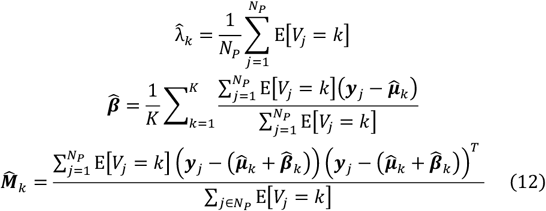

The E-step and M-step are iteratively applied till convergence. In certain cases, we can choose to fix 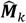 as 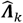 to make the shape of cell clusters in the perturbation group similar to those in the control group.

## 3 Results

### 3.1 Performance of scDecouple on synthetic data

We conducted a series of simulation experiments to demonstrate the performance of scDecouple in decoupling cellular response from infected proportion bias. We first generated a synthetic dataset. We randomly generated 1,000 two-dimensional NT cells as the control group. These cells followed a two-cluster GMM (Fig. 3A), whose bimodality concentrates on PC 1. We used the same probability model to randomly generate the initial states of cells in the perturbation group, and added perturbation effects on these cells with the assumption that all cells exhibit similar responses to the perturbation in the PC space. According to Eq. 3, the infected proportion bias of any PC in the scenario of two clusters is

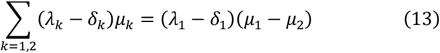

**Fig. 3.**
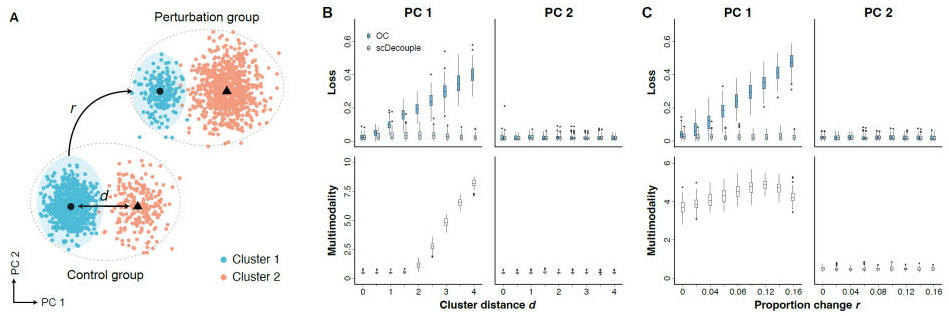
Schematic diagram of synthetic data and performance of different methods on synthetic data. **(A)** The illustration of synthetic data.**(B)** Performance of different methods on synthetic data with dif-ferent cluster distances *d*. **(C)** Performance of different methods on syn-thetic data with different ratio changes *r*.

The bias was linearly correlated with two components: the distance *d* between two clusters and ratio change *r* of cluster 1 between two groups:

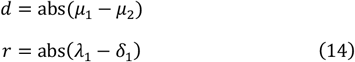

By varying the values of *d* and *r*, we systematically controlled the magnitude of infected proportion bias. For each parameter combination, we calculated OC as the results of naive analysis and applied scDecouple to estimate real cellular response. We compared the results of these methods with ground truth. We also evaluated the effectiveness of the Hartigan’s dip test as the metric for assessing the inherent multimodality of the control group.

We first fixed the ratio change *r* = 0.1 and varied the cluster distance *d*(Fig. 3B), and then fixed *d* = 3 and varied *r* (Fig. 3C). We conducted 100 simulations for each parameter setting. When *d* or *r* was small, OC and the scDecouple results both exhibited low losses on PC 1 (Fig. 3BC). However, as the increase of data multimodality or ratio change, estimation by OC showed higher losses, whereas scDecouple continued to perform well. In the PC 2 dimension where the data distribution is unimodal, both methods showed similar results (Fig. 3B-C). Besides, our results demonstrated that the Hartigan’s dip test in scDecouple exhibited high sensitivity to changes in multimodality (Fig. 3B-C).

The results revealed that when facing cellular heterogeneity and uneven gRNA sampling across different cell clusters, OC can hardly be regarded as cellular response as it is affected by infected proportion bias. In contrast, scDecouple decouples these intertwined factors and consistently produces low-error estimation outcomes, irrespective of the degree of infected proportion bias.

### 3.2 Performance of scDecouple on simulated biologically-derived genome-wide data

We conducted simulations using real scCRISPR-seq data to further evaluate the performance of scDecouple. We collected a scCRISPR-seq dataset from a genome-wide Perturb-seq study^15^ and selected the experimental data from two distinct cell lines, K562 and RPE1. We mixed these data to simulate two cell clusters in a single experiment. We found that the cell number infected by different gRNAs varied in both cell lines (Fig. 4A). The correlation of the cell number infected by the same gRNA between different cell lines is notably low (Pearson’s correlation coefficient = 0.155, Fig. 4B), suggesting significant infected proportion bias between cell clusters.

**Fig. 4.**
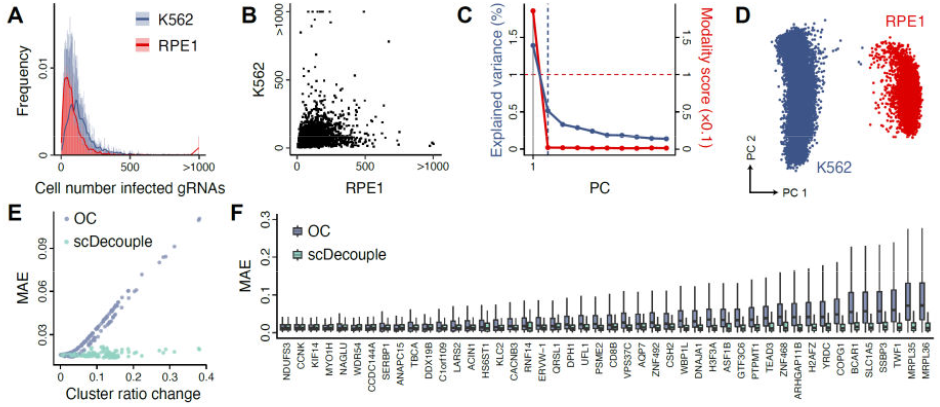
Performance of different methods on simulated genome-scale scCRISPR-seq data. **(A)** The distribution of the cell number infected by different gRNAs in different cell lines. Solid lines show the density of distributions. **(B)** The cell number infected by the same gRNA between different cell lines. **(C)** The explained variance and multimodality score of each PC. Dashed lines represent thresholds for identifying high variance or multimodality. **(D)** The distribution of cells in PC 1 and PC 2. **(E)** Cluster ratio changes and mean absolute errors (MAEs) of cell response estimations on PC1. Each Dot represents a gRNA. **(F)** MAEs of cell response estimations in gene expression profiles. One sample represents a gene in the gene expression profile. The x-axis represents gRNA targets, sorted by cluster ratio changes.

We applied scDecouple on the simulated data. We found that PC 1 showed both the highest explained variance and the highest multimodality score among all PCs (Fig. 4C-D). Thus, we performed decoupling on PC 1 with the cell cluster number *K* = 2 and regarded OC as the cellular response for all other PCs. For the perturbation group, we selected 168 editing loci which were designed in both cell line groups and showed similar responses in both cell types (the difference of cellular response on PC 1 less than 5). We took the mean responses (log FC) of the two cell lines on PC 1 as true cellular response and used the mean absolute error (MAE) to assess the accuracy of cell response estimations. We found that scDecouple can better estimate true cellular response than OC, especially when cells infected by a gRNA displayed a substantial cluster ratio change (Fig. 4E). We then randomly selected several gRNAs and generated the cellular response of these gRNAs from the PC space to gene expression. We found that responses calculated by scDecouple showed lower MAEs than OC, especially when ratio changes were large (Fig. 4F). In general, the results showed that scDecouple revealed more accurate cellular responses and reduced the error introduced by infected proportion bias.

### Applying scDecouple on real scCRISPR-seq data

We applied scDecouple to a real scCRISPR-seq dataset targeting T cell receptor (TCR) related genes in human Jurkat cells^5^. Each gene was targeted by three gRNAs. We selected 22 target genes each with over 60 infected cells. We considered the cells affected by gRNAs targeting the same gene as one perturbation group. We then got 22 perturbation groups along with NT gRNAs as the control group for downstream analysis. We performed data preprocessing on the data and then transformed the data into the PC space of the control group based on the top 700 highly variable genes. We identified three PCs with both high explained variances and high multimodality scores (Fig. 5A). We performed decoupling on these three PCs with the cell cluster number K = 2. We found that the estimated cell cluster proportion for each target varied (Fig. 5B), and the inferred clusters showed distinct differences in the PC space (Fig. 5C). We generated the cellular response of all targets from the PC space to gene expression obtained the average response of a gene by averaging the response of all gRNAs targeting this gene. We visualized the responses of TCR pathway signature genes defined in the original publication^5^ (Fig. 5D). The results showed that knockouts of some genes caused pathway activation, while others led to pathway inhibition.

**Fig. 5.**
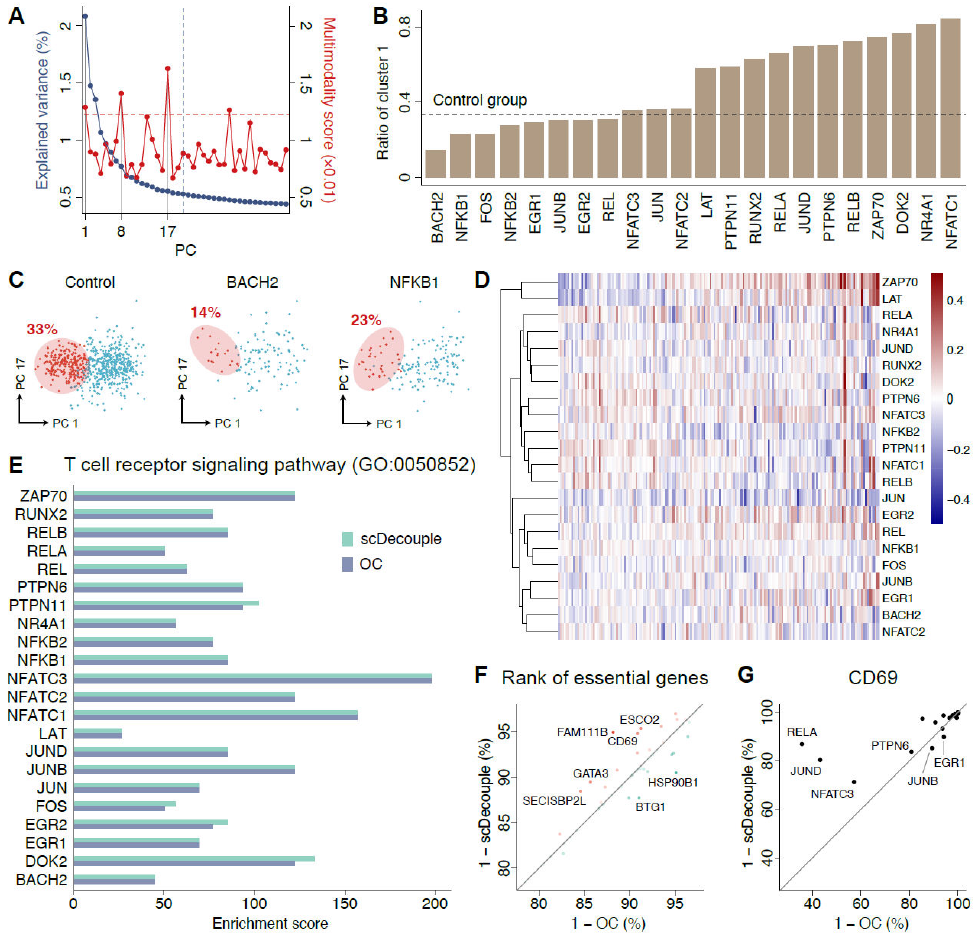
The performance of scDecouple on real scCRISPR-seq data. The explained variance and multimodality score of each PC. Dashed lines represent thresholds for identifying high variance or multimodality. The ratios of cluster 1 across all targets. **(C)** The distribution of cells in PC 1 and PC 17. **(D)** Cellular response estimations of all targets in the expression of all TCR pathway-related signature genes. **(E)** The enrichment scores of differential genes in the TCR signaling pathway for each target. **(F)** Average gene ranking of top 60 genes from the TCR pathway signature genes across all gRNAs. Each dot represents one signature gene. **(G)** Ranking of CD69 in the absolute values of cellular responses calculated by OC or scDecouple. Each dot represents one target.

We then compared the cellular response estimated by OC and scDecouple. We identified genes with the top 1500 absolute values of cellular responses as differential genes for the OC and scDecouple results, respectively, and then calculated the enrichment score of the TCR signaling pathway for each target by the Fisher’s exact test (Fig. 5E). The results showed that scDecouple achieved similar or higher enrichment scores than OC.

We further investigated whether scDecouple helps biological discovery. We selected the top 60 genes from the TCR pathway signature genes according to the original study^5^, and calculated the average rank of these genes in the absolute values of cellular responses calculated by OC and scDecouple, respectively. We found that these genes showed higher rank in scDecouple results (Fig. 5F). We specifically examined CD69, a widely recognized early activation marker of the TCR pathway^21,22^. The results demonstrated that CD69 rankings were greatly improved for most targets (Fig. 5G). The most significant improvement showed in the perturbation group targeting RELA which was reported to be highly associate with CD69^24–26^. All the results indicated that scDecouple can benefit the analysis of real scCRISPR-seq data with more precise gene ranking and more accurate gene regulation identification.

## 4 Discussion

We initially formalized the process of cellular response estimation in scCRISPR-seq and derived mathematical equations to deduce estimation bias induced by infected proportion bias. To reduce the influence of infected proportion bias, we introduced scDecouple, a method aimed at unraveling observed changes in scCRISPR-seq data. Our approach focuses on decoupling cellular response from infected proportion bias through maximum likelihood estimation. We validated the efficacy of scDecouple through a series of simulations and its application to a real dataset. scDecouple yielded more enriched pathways and improved the ranking of perturbation-related genes. The good performance of scDecouple is mainly attributed to its estimation of the actual proportions of clusters for infected cells in perturbation groups.

With the development of technology, scCRISPR-seq with more complex gRNA libraries will be developed and applied to populations of more heterogeneous cells. As a result, reducing infected proportion bias will be more important and necessary in the analysis of scCRISPR-seq. Besides, Double-strand breaking induced by CRISPR knockout may cause cell state arrest, which will introduce additional bias to cell cluster proportion especially in studies related to cell cycle, senescence or aging. scDecouple can help to discard these impacts and focus on the cellular responses we concern about.

We have built an R package for scDecouple, offering a comprehensive suite of functionalities for streamlined scCRISPR-seq data analysis. The package comes equipped with an integrated one-click data preprocessing pipeline, infected proportion bias quantification, cellular response estimation, and downstream analytical tools. Additionally, the library provides many visualization options, including PC plots, PC variance and multimodality visualizations, cellular response heatmaps, Gene Ontology (GO) enrichment plots, etc. Moreover, the library supports both step-by-step execution and parameter customization to meet the diverse needs.

As the first method designed to handle infected proportion bias in scCRISPR-seq, scDecouple can be further optimized. Currently, scDecouple operates under three key assumptions: cells following GMM within the PC space, same cell clusters between NT and perturbation groups, and similar cellular responses across cell clusters. The first assumption holds well in certain cell lines and tissues, but it might not remain in intricate systems or disease tissues, such as tumors. We can expand scDecouple by considering alternative data representation methods or statistical models to handle more complex tissues and systems. The second assumption might not hold true in extreme cases, such as when some rare cell types are completely absent from infection. In such a situation, the number of clusters in the control and perturbation group could differ. We plan to incorporate additional information to address this problem, such as including marker genes of cell clusters to link clusters in the perturbation group to those in the control group. The third assumption guarantees that scDecouple is suitable for cells with moderate heterogeneity, which is validated in this study by applying scDecouple on real scCRISPR-seq data. In the future, we intend to integrate more intricate models, such as deep learning, with scDecouple to handle more complex experimental data.

## Funding

This work is supported in part by National Key R&D Program of China (2021YFF1200900, 2019YFA0904402, 2019YFA0906700), National Natural Science Foundation of China (62250005, 61721003, 32171448, and 62103227), Tsinghua University Initiative Scientific Research Program (2022Z11QYJ032, 2021Z11JCQ020), Beijing Natural Science Foundation (Z210010), Tsinghua University Spring Breeze Fund (2020Z99CFG006), and the AIR@InnoHK administered by Innovation and Technology Commission of Hong Kong.

### Data and code availability

We only use public datasets in this study. The K562 and RPE1 datasets employed for simulations were obtained through accession code GSE92872. The Jurkat cell dataset was downloaded from the website https://gwps.wi.mit.edu.

All codes of this study, including the R package of scDecouple, simulation experiments, and real scCRISPR-seq data analysis, are available on GitHub via the following link: https://github.com/MengQiuchen/scDecouple.

